# StrainVis: interactive visual strain-level analysis of microbiome data

**DOI:** 10.64898/2026.03.11.711087

**Authors:** Inbal Paz, Ruth E. Ley, Hagay Enav

## Abstract

**Background:** Microbiomes contain multiple conspecific strains whose genomic differences arise from both single nucleotide variants (SNVs) and structural variation (insertions, deletions, recombination). Recently, computational tools to assess strain-level differences became available, based either on average nucleotide identity (ANI) or on the average pairwise synteny (APSS) of strains, which are sensitive, respectively, to either SNVs or to structural variation. However, strain-level analyses remain technically challenging and fragmented across approaches and combining these complementary signals typically requires substantial bioinformatic expertise.

**Results:** Here we present *StrainVis*, a web-based analysis and visualization platform that integrates outputs from both ANI- and APSS-based strain tracking tools to enable unified, interactive exploration of within-species diversity. StrainVis allows users to perform per-species and multi-species comparisons, incorporate metadata and gene annotations, and generate statistical summaries and publication-ready figures without programming.

**Conclusions:** By lowering technical barriers and enabling joint interpretation of sequence and structural variation, *StrainVis* makes advanced strain-level microbiome analysis accessible to a broader community and facilitates discovery of evolutionary patterns that would be missed by single-method approaches alone.

## Introduction

Microbiomes are collections of microbial species that often include a high number of different strains. Strains of the same species, termed conspecific strains, differ in their genotypes due to accumulation of SNVs (Single Nucleotide Variants) or due to processes such as deletions, insertions, recombination events and horizontal gene transfers, which result in genomic structural differences. Both SNVs and structural genomic variation were previously shown to have the potential of modifying the microbial phenotype [1, 2] and in the context of the human microbiome, could have a different effect on the host [3–9].

In spite of the contribution of genomic structural variation to the generation of within-species strain diversity, most tools to track and compare conspecific strains are based on differences in SNV between genomes (reviewed in [10]) and show lower sensitivity to genomic structural variation. This, in turn, could lead to underestimation of the strain-diversity in species with high recombination rates. Moreover, SNV-based tools could overestimate genomic diversity due to technical or biological factors, for example, sequencing errors or the presence of hypermutators with increased point mutation rates [11, 12].

The relative shortcomings of the SNV-based tools motivated us recently to develop a new metric for quantifying the similarity between strains based on differences in microsynteny. *SynTracker* [13], a computational tool for tracking and analyzing conspecific strains based on genome microsynteny - i.e., the conservation in the order of short k-mers along two chromosomes. *SynTracker* quantifies the similarity between different strains within a species based on structural genomic variation information contained in genomic and metagenomic datasets. In a benchmarking assay, *SynTracker* outperformed other SNV-based strain tracking tools. Moreover, by combining synteny- and SNV-based strain tracking, we were able to define the per-species evolutionary mode, i.e., identify species evolving by preferential accumulation of point mutations or genomic structural variation.

Most computational pipelines and software in the microbiome field, including *SynTracker*, output tables which should be further processed to reach biological insights. This requirement limits the use of these tools by making downstream analyses accessible only to users with bioinformatic and programmatic skills and background. These hurdles slow down the assimilation of strain-level analyses in the microbiome field particularly by new users. A good solution is to deploy graphical user interface (GUI), many of which were developed in the microbiome field to facilitate the use of computational methods by experimentalists and bioinformaticians alike. For example, the *QIIME2* platform [14] includes a GUI feature, allowing visualization of the platform’s taxonomic analyses. *Anvi’o* [15] is a software providing a GUI based interactive platform for the analysis and visualization of microbiome data. *CViewer* [16] is a java-based statistical framework, allowing exploration and analysis of metagenomic data without need for specialized skills. Another approach is presented by *VAMPS* [17] and *WHAM! [18]*, two web based tools for the visualization of microbial community data. To date, there is no such GUI for strain-level analyses.

Here, we developed *StrainVIs*, a web-based GUI that provides accessible, advanced strain-level analysis and visualization capabilities for both experimentalists and bioinformaticians. *StrainVis* enables interactive analysis and visualization of outputs from *SynTracker* and ANI-based strain tracking tools. *StrainVis* takes as input tables produced by strain analysis tools and supports two main types of strain-focused analyses: per-species analyses, which enable in-depth investigation of conspecific strains of a given species, and multi-species analyses, which compare strain similarities across multiple species. In addition, *StrainVis* enables further investigation of within-species diversity, by accepting metadata and gene-annotation files. Overall, *StrainVis* facilitates high-resolution microbiome analysis at the strain level by enabling strain tracking, statistical testing, and the generation of publication-quality visualizations.

*StrainVis* is an open-source tool, available for download under: https://github.com/leylabmpi/StrainVis/

## Methods & Implementation

### Tool design

*StrainVis* was designed to (A) allow non-bioinformatician scientists to perform in-depth strain-resolved analyses of complex microbiome data, and (B) encourage standardized analysis workflow by both bioinformaticians and experimentalists. StrainVis is not a standalone tool, but instead designed as an extension of existing strain tracking tools. Specifically, StrainVis relies on the output of *SynTracker* [13], a tool to track strains using genome microsynteny and on the output of ANI-based tools, for example *inStrain* [19]. These two types of outputs highlight two complementary modes for estimating strain relatedness - with *SynTracker* the measure for strain similarity is the Average Pairwise Synteny Score (APSS), which is based on structural genomic variation and not on SNVs. On the other hand, ANI-based tools rely mostly on SNVs to determine strain similarities, while being less sensitive to structural genomic variation. We previously showed that combining these two approaches could shed light on the preferential mode in which different species evolve [13].

As *StrainVis* is intended to be used by both experimentalists and bioinformaticians, we decided that a straightforward approach should be a leading concept in its design. Our tool is designed as a python based web-application, however, all analyses and plots are generated locally on the user’s machine, allowing greater independence while minimizing potential data transfer and sharing issues.

### Technical overview (visualization and computational tools)

The *StrainVis* application is available for download as a single archived file. The tool is built using ***Panel*** *[20]*, a Powerful interactive Data Exploration & Web App Framework for Python. The StrainVis web-application is served in the browser using the ***Bokeh*** [21] server. A single command in the machine’s terminal launches the Bokeh server and opens the application in a web-browser window under a dedicated URL. As long as the Bokeh server is not interrupted, the StrainVis application can be accessed using the same URL. Our tool allows multiple simultaneous instances of the program to run at the same time if the analysis of multiple datasets in parallel is required. The Bokeh server, serving the StrainVis application, can also be executed on a remote machine, with the application being accessible from the local browser.

### Data input and output

StrainVis accepts four types of input data:

I. A single file, containing the output table of *SynTracker*. Each line of this table specifies the species being compared, the region of the genome being tracked (denoted by its coordinates along the reference genome), the metagenomic samples (or isolates) being compared and the corresponding Synteny score [13]. This table can include multiple species.
II. A single file, containing the ANI values. Each line of this table contains the Species being compared, the compared sample pair and the ANI computed for the species pairwise comparison. This table can include multiple species.
III. An optional table containing the metadata for each sample (or genome, if microbial isolates are analyzed). The metadata file, in a tab delimited format, can contain any type of per-sample (or per-genome) information, relevant for the downstream analysis. This information can be discrete, categorical or continuous. There is no limitation for the number of metadata fields.
IV. An optional gene annotation file, per each reference genome analyzed.

Following the launch of StrainVis, the input file selection screen appears in the browser and allows the user to specify whether they are interested in Synteny-based analysis, ANI-based analysis, or both, and to upload the relevant input files.

While all StrainVis analyses and visualizations are displayed on-screen, analysis results and data tables could also be exported as table files. All visualizations can be exported in one of the following image formats: png, pdf, svg and eps.

### Data analysis and visualization

StrainVis has two modes of data analysis and Visualization: per-species and multi-species (Fig. 1). In the per-species mode, the user selects the species of interest from a drop-down menu, containing all species identified in the input files. If the user uploaded both *SynTracker* and ANI files three types of analyses (Synteny-based, ANI-based and combined) are presented using separate tabs.

**Figure 1:**
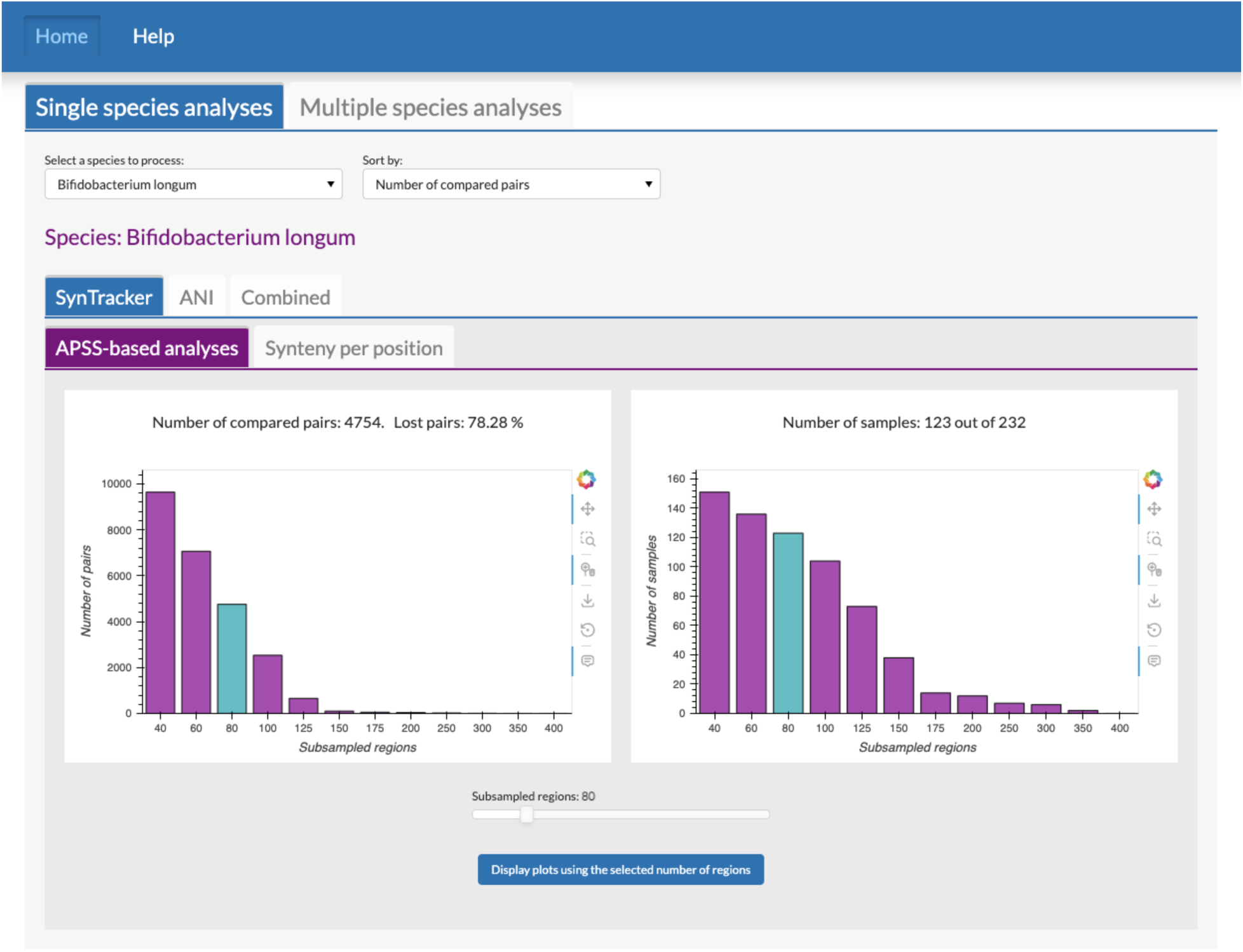
View of the initial interactive screen of StrainVis. The user selects either single species or multi species analyses (upper tabs) and whether the analyses are based on APSS, ANI or both (second row tabs). In the APSS-based analyses the user selects interactively, using a slider, the number of genomic regions used per pairwise-comparison, and is informed on the number of samples (right) and pairwise comparisons (left) included in downstream analyses.

Interactive synteny-based analyses included in the per species mode are (1) analyses based on *Average Pairwise Synteny Scores* (APSS), calculated using ***n*** subsampled regions per-pairwise comparison (the number of regions is determined by the user and could be changed interactively), and (2) visualization of the synteny scores along each position in the species reference genome. APSS-based visualizations include **initial bar-plots** (Fig. 1), allowing the user to modify the number of sub-sampled regions while being informed about the influence of the subsampling on the number of samples and sample-pairs being analyzed. The **APSS-distribution plot** is a jittered scatter (or a box) plot, showing the distribution of APSS of the species, and optionally for different metadata categories. The **clustered heatmap** displays sample clustering based on APSS values and can optionally include an additional row corresponding to any metadata field. The **Network plot** is a per-species network, in which the nodes correspond to the samples and the edges connect sample pairs with APSS score above a user-determined threshold. The network is displayed in a force-directed layout, i.e., nodes are clustered, using the Fruchterman Reingold algorithm, to visualize the similarities between groups of samples, based on their APSS values. Additionally, both the nodes and the edges of the network could be colored by any metadata field found in the metadata file. The parameters for all visualizations could be controlled by the user, using the StrainVis GUI. The **Synteny per-position** plot is composed of several sub-plots, displaying the individual synteny scores and the population average, along each position of the reference genome. In addition, hyper-conserved (10% of the regions with the highest score, that also appear in at least 50% of the sample pairs) and hyper-variable regions (10% of the regions with the lowest synteny scores that were identified in at least 10% of the samples) are highlighted. Each of the sub-plots can be interactively set as shown or hidden by the user. The presented information could be filtered by different metadata fields. Gene annotation data could also be displayed, once an annotation file for the reference genome is provided.

Similarly, the ANI-based per-species visualizations include the distribution plot, the clustered heatmap and the network graph, all based on the pairwise ANI values.

The combined mode includes a scatter plot, presenting the ANI values vs. the APSS and the Spearman correlation between them, with an optional coloring of the data points according to different metadata categories.

The multi-species mode is activated when the user provides an input table containing the analysis of more than a single species. In this analysis mode, the user may include all species or select a subset and display the distribution of strain-similarities among them. Optionally, the user can compare different metadata categories, while performing all statistical tests in real-time. When the input is composed of both *SynTracker* and ANI results, two separate tabs are displayed, one for APSS-based and the other for ANI-based analyses.

## Results

To showcase the functionality of *StrainVis*, we first applied it to a previously published human-gut metagenomic dataset composed of samples from infants and their mothers, collected over 3 countries: Gabon, Vietnam and Germany [22]. This dataset consists of 1133 samples assembled into sample-specific MAGs (metagenomic assembled genomes). We grouped MAGs into species clusters, using an ANI cutoff of 95% and identified a Species Representative Genome (SRG) for each species-cluster as described previously [22]. The SRGs were used as reference genome for both the synteny-based and ANI-based strain analyses (using *SynTracker* and *inStrain [19]*, respectively).

In this analysis, we decided to investigate a subset of the most prevalent species in our dataset, i.e., the 10 species with the highest number of comparisons between samples. We first used the Multi-species analysis feature of StrainVis, and continued with an in-depth, per-species analysis of two species of interest.

## Multi-species analyses

In the multi species analysis the pairwise strain similarities, either APSS or ANI, for all species are plotted together. The user can further divide the within-species groups to subgroups, according to different metadata categories. We plotted the strain similarity values for the set of species (using 80 regions per pairwise comparison, for the synteny based analysis) focusing on the following metadata fields: (a) Country, (b) Province and (c) infant/adult. Overall, both analyses show similar trends: most of the species show significantly higher strain-level similarity in sample-pairs coming from the same province compared to those coming from different provinces, and those coming from the same country compared to those coming from different countries (8 and 9 significantly different species for the synteny-based and ANI-based same/different country comparisons, Fig. 2a, and 7 and 8 species with significant differences for the same/different province, Fig S1).

**Figure 2:**
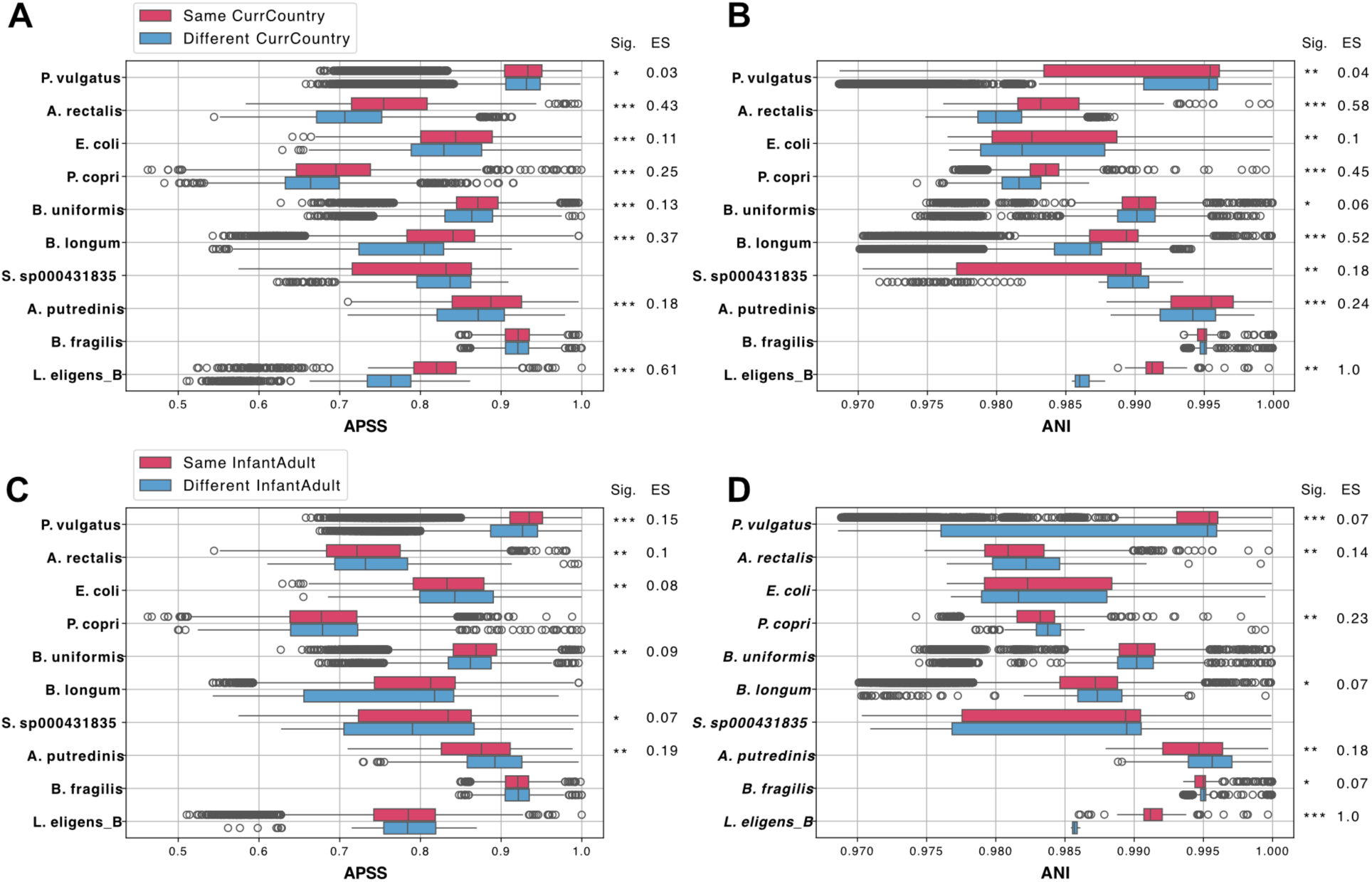
Multi-species analyses of strain-similarities of the most abundant 10 species in the human gut metagenomes collected in Gabon, Germany and Vietnam. (A) Synteny-based (left) and ANI-based (right) comparisons, with internal within-species classification to sample pairs in which the hosts reside currently in the same country (i.e, Same_CurrCountry) or not. (B) Synteny and ANI based comparisons, with pairwise comparisons classified as “Same Infant/adult”, i.e., both samples in a pair obtained from an infant, or both from an adult host, or classified as "Different infant/Adult”, meaning a pairwise comparison in which one sample was obtained from an adult subject, while the other was obtained from an infant. Stars correspond to Benjamini–Hochberg-corrected P values (two-sided Mann-Whitney U rank test). *q < 5 × 10^−2^, **q < 5 × 10^−5^ and ***q < 5 × 10^−10^. Effect size calculated using the Rank-Biserial correlation method.

We next explored the strain similarities of the set of species, when grouping pairwise comparisons into same or different “adult/infant” categories (i.e., adult-adult and infant-infant comparisons in the “same” group, and infant-adult comparisons in the "different" group). We observed 7 species with a significant difference between the groups in the ANI analysis and 6 species with a significant difference in the synteny-based analysis (Fig 2b). However, in many of the species the average strain similarity was higher in the adult-vs-infant comparisons, compared to infant-infant and adult-adult groups. Therefore, we suggest that geography plays a bigger role in determining the within-species similarity, compared to the formation of distinct clades, typical to the adult or infant gut microbiome.

## Per species analyses

Next, we used the per-species analysis mode to conduct a high-resolution study of two distinct species from our dataset: *B.longum* and *L.eligens*. We performed the synteny based analysis using a selection of 80 regions per pairwise comparison for *B.longum*, and 60 regions for *L.eligens,* resulting in 4754 pairwise comparison for *B.longum* and 2440 for *L.eligens*. In the ANI-based analysis 10318 and 141 sample pairs were included, respectively.

### APSS and ANI distributions

Within species genomic diversity results from the accumulation of genomic structural differences (reflected in the APSS), SNVs (reflected in the ANI) or both together. We previously showed that some species evolve by preferential accumulation of one type of genomic differences, while others accumulate both at a similar rate [13]. To investigate the mode in which the two species evolve, we plotted the pairwise ANI against the APSS values of each species (Fig 3). This analysis revealed that while the Spearman rank correlation of the two strain-similarity measures for *B.longum* is relatively high (*r = 0.82*), the two measures correlate poorly for *L.eligens* (*r = 0.46*), suggesting that in this species, the different types of genomic variation do not accumulate at similar rates in different strains.

**Figure 3:**
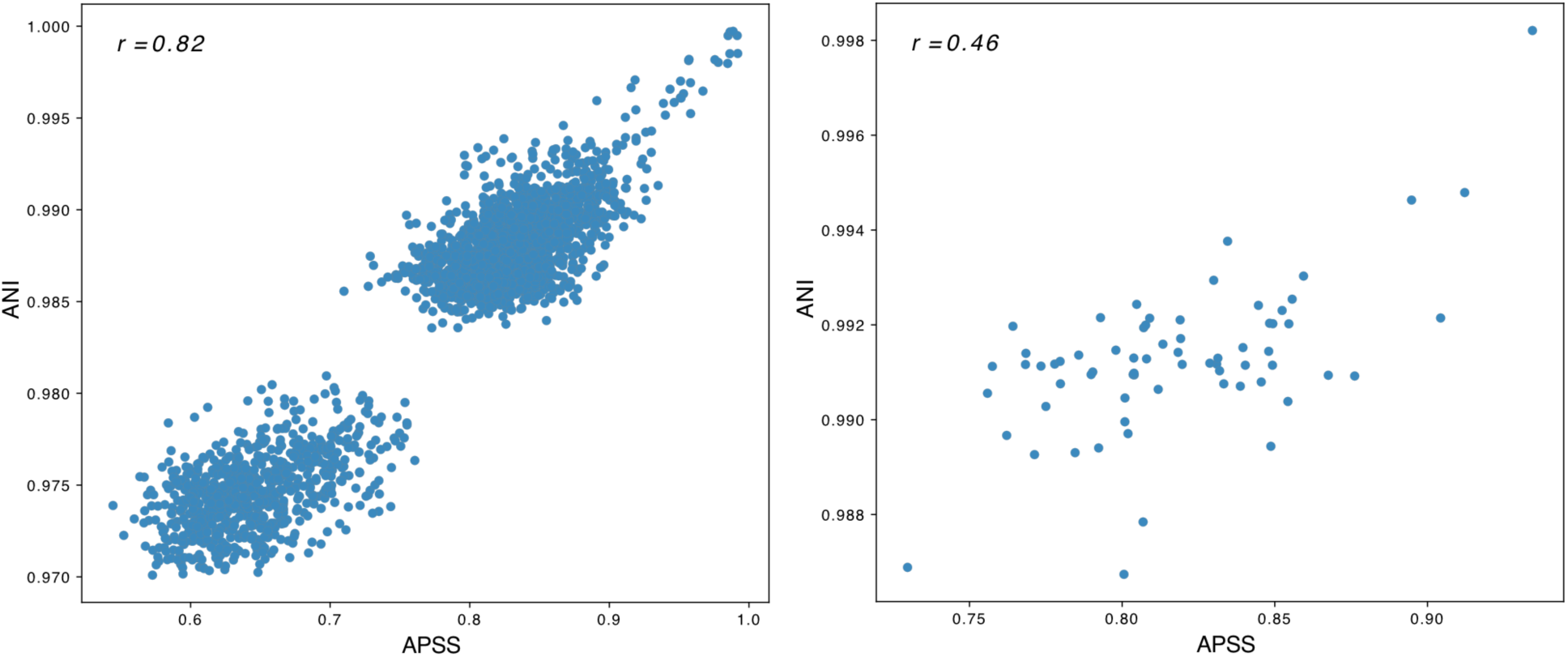
SNP-based strain similarities (i.e., ANI), plotted against the synteny-based strain similarities (i.e, APSS) of members of two species. *B.longum* (left) and *L.eligens* (right). Each datapoint represents a pairwise comparison of the members of each of the species, in two distinct metagenomic samples. Sample pairs included are only those detected by both tools: 2640 *B.longum* pairs and 68 *L.eligens* pairs.

We next used the boxplot tool to visualize the strain similarities for *B.longum*. We observed that a striking difference exists between pairs coming from the same village or town and pairs coming from different villages (p =< 7.9 x 10^-51^, effect-size = 0.35 for the APSS-based comparison and р =< 3.8 x 10^-235^, effect-size = 0.5 for the ANI-based comparison) (Fig 4a, 4b). This finding supports the possibility of transmission of this species within geographically distinct populations. We created a jitterplot for the *B.lungom* pairwise comparisons, demonstrating a clear bimodal distribution, both for the APSS and ANI based strain similarities (Fig 4c, 4d). This distribution could point to the existence of few *B.longum* “clades” with high within-clade similarity, while the pairwise comparisons with lower genomic similarity reflect comparisons of members of different clades.

**Figure 4:**
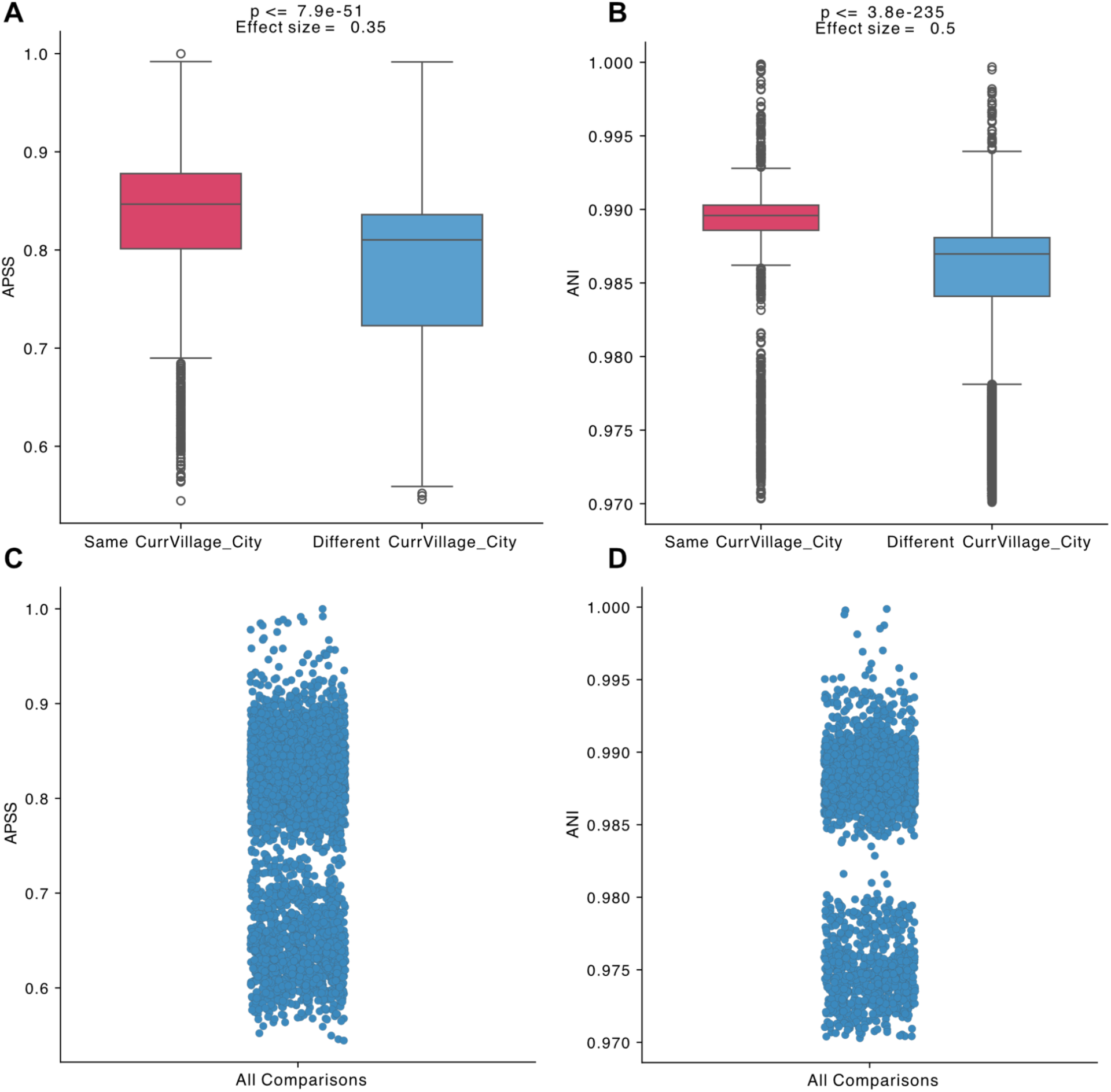
Pairwise strain similarities of *B.longum*. (A) Synteny-based and (B) ANI-based strain comparisons, grouped into pairs coming from the same village or town, or those coming from different villages/towns.(C) Jitter scatterplot showing the bimodal distribution of APSS and (D) ANI pairwise comparisons. Between groups p-values are the result of the Mann-Whitney U test, effect size calculated using the Rank-Biserial correlation method.

### Clustered Heatmaps and network

To visually explore our hypothesis regarding the existence of a few distinct clades of *B.longum* we used the clustered heatmap feature of *StrainVis*. We created an APSS and ANI based symmetric heatmaps, clustered using the “correlation” option. To explore potential correspondence between strain relatedness and the location of the host, we colored the sample nodes by the host’s country of residence. Overall, both the APSS-based and ANI-based heatmap were in agreement and reflected the existence of two major sample-clusters, differing in size. While in both heatmaps the smaller cluster is composed of samples from different countries, without a clear internal organisation, the larger cluster, containing the majority of the samples, has an internal organisation reflecting the geographic location of the hosts (Fig 5a, 5b).

**Figure 5:**
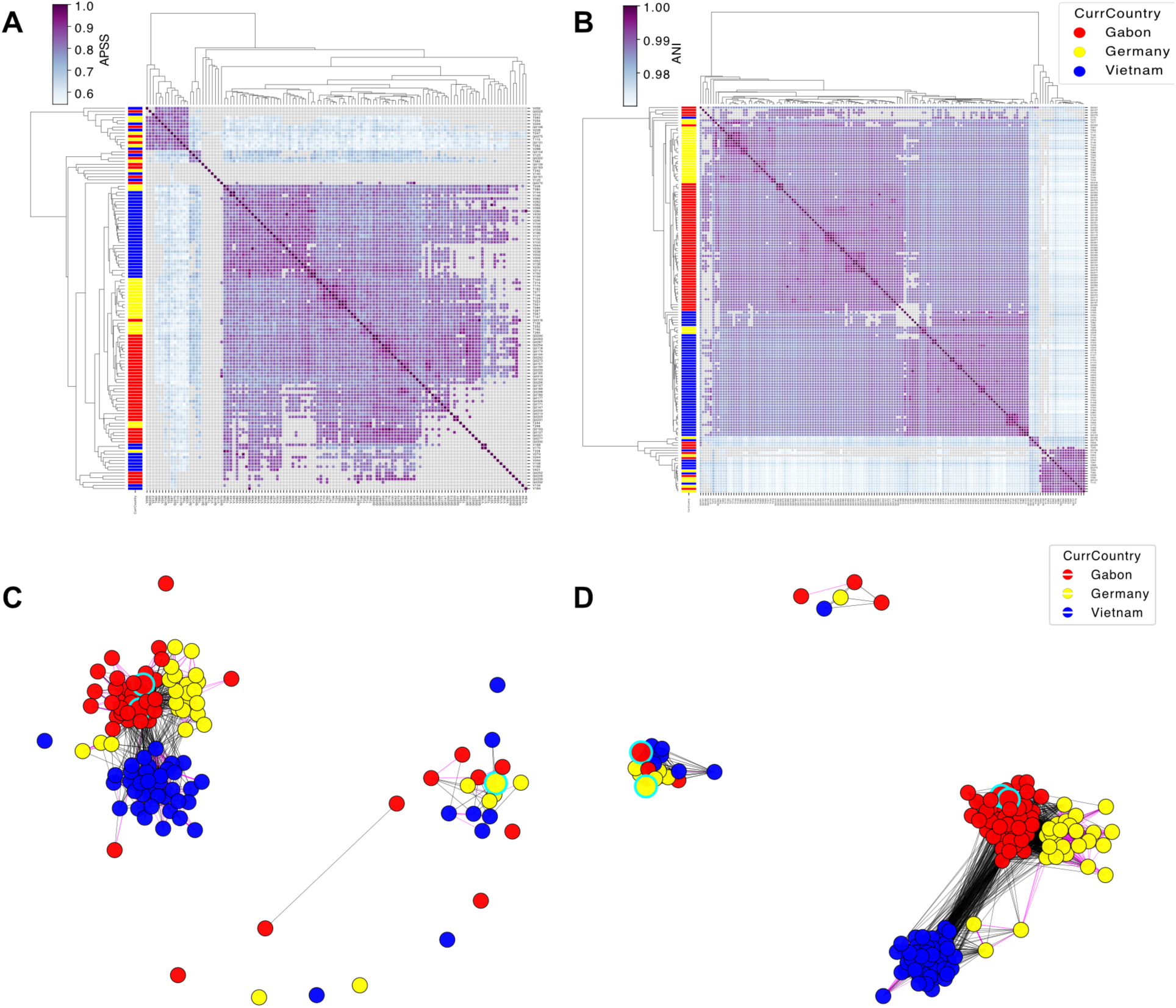
Visualization of the within species genomic similarities of B.longum conspecific strains. (A) APSS based strain similarity heatmap. Left side color annotations represent countries: red - Gabon, yellow - Germany, blue - Vietnam. (B) ANI based heatmap. (C) APSS based strain similarity clustered network. Node color represents the host’s country of residence, edge color represents whether two hosts reside in the same province (pink) or in different provinces (black). Samples obtained from adult hosts are highlighted, using the metadata feature, and circled in cyan color. (D) same as in C, using ANI to estimate strain relatedness.

To further explore the population structure of *B.longum*, and to verify our findings, we used the “clustered network” feature. In this network-visualization members of the species are represented as nodes. Two samples with a strain similarity that exceeds a determined threshold (in this analysis, 0.85 for the APSS-based network and 0.988 for the ANI-based network) are connected with an edge and weighted according to the similarity between the two members of the species. The nodes are then clustered based on their edge weights, to allow visual inspection of the genomic landscape of this species. In these networks we used 400 and 300 iterations for the APSS and ANI based networks, respectively. We colored the nodes according to the host’s country of residence and colored edges to reflect if the two connected samples originate in the same province or not. Since *B.longum* is mostly identified in infants, we highlighted nodes representing adult hosts (3 in the *SynTracker* analysis and 4 in the ANI based analysis), to understand if adult-strains are clustered together. Examination of both networks reveals the existence of two completely distinct clusters. The bigger cluster is composed of three major sub-clusters, each mainly composed of strains originating in a single country. The smaller cluster is composed of samples from all three countries. These findings further support our observation that the *B.longum* global population is composed of few distinct “clades” with high within-clade genomic similarity but with low between-clade similarity. Moreover, the networks show that the adult samples are located in different clusters, therefore the genomic differences are not a result of the existence of adult/infant genotypes (Fig 5c, 5d).

### Per-position synteny plots

Genomic differences are not equally distributed along genomes, due to the existence of “hot-spots” of recombination, horizontal gene transfer and prophage integration sites [23] and as selective forces act differently on different genes. In order to explore regions of higher divergency or conservation, we created the per-position synteny plot feature, where the average and individual pairwise synteny scores are plotted per each 5 kbp window along the reference genome. The synteny per-position plot can be overlaid with gene annotations, providing per-position diversity measures in the context of regional gene content.

Here, to showcase the abilities of this feature, we explore in detail contig 630 from the *B.longum* reference genome. When inspecting the per-position diversity of the 280 kbp long contig, based on the entire sample cohort, *StrainVis* defined four 5 kbp regions as hyper-variable, and 3 regions as hyper-conserved (Fig 6a). Interestingly, two of the hyper-conserved regions are adjacent (positions 205,000-215,000). Hyper-conserved and hyper-variable regions differed markedly in gene content: hyper-conserved regions were enriched for genes encoding non-hypothetical proteins (*p* ≤ 0.0032), whereas hyper-variable regions were enriched for genes encoding hypothetical proteins (*p* ≤ 6.1 × 10⁻⁴; hypergeometric test; Fig 6b), suggesting that that hyper-variable genes, at the strain level, are less represented across taxa and in databases used for annotating genes.

**Figure 6:**
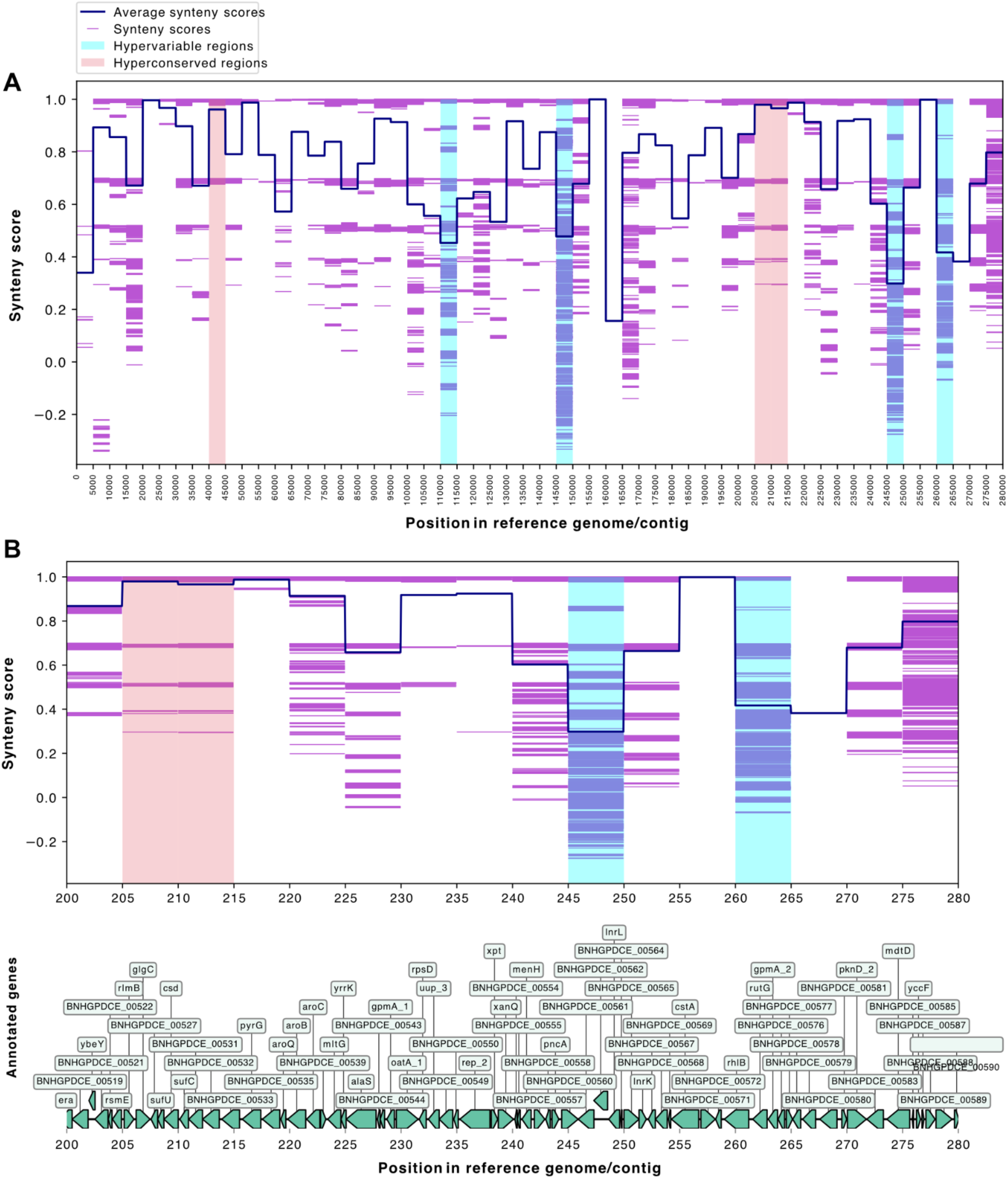
Synteny per-position plot of contig 630 of the *B.longum* reference genome, for the entire host population. (A) Synteny per position of the entire contig, (B) zoom-in view of positions 200,000-280,000, overlaid with predicted gene annotations. Horizontal purple lines represent individual pairwise synteny scores for each region, the dark blue line represents the average synteny score for all pairwise comparisons, in each genomic position. Vertical blue highlighting represents hyper-variable regions, salmon vertical highlighting represents hyper-conserved regions.

We further used the “filter plot by metadata” feature to investigate whether local populations, i.e., residing in each of the countries, show different patterns of conservation and variation in this contig. We observed that while the hyper-variable regions in each of the populations partially overlap, large differences exist in the hyper-conserved regions identified in each sub-populations. We hypothesise that different selective forces in each of the subpopulations purge genomic variations in different genes, as different functions are preserved through purifying selection, in each local population (Fig S2).

## Conclusions

We present here *StrainVis*, a comprehensive GUI-based computational tool for advanced downstream analyses of microbiome data at the sub-species level. *StrainVis* allows the entire scientific community, with an emphasis on non-bioinformaticians, to execute, in a time efficient manner, high-resolution studies of microbial dataset, while incorporating matching environmental and medical metadata. *StrainVis* can use the output of both *SynTracker* and of ANI-based strain tracking tools, to allow strain analysis using different types of genome variation.

## Author Contributions

REL conceived the project. IP, HE and REL designed the tool. IP did the programming, graphical design and wrote the manual. HE performed analysis and wrote the manuscript with help from all authors.

## Acknowledgements

We thank members of the Department of Microbiome Science for their constructive feedback. This work was supported by the Max Planck Society. The authors declare no competing interests.

## Declarations

Ethics approval and consent to participate: not applicable. Consent for publication: not applicable.

## Availability of data and material

*StrainVis* is publicly available - https://github.com/leylabmpi/StrainVis.

Input tables and files are publicly available [24].

## Extended methods

### Python packages used in the visualization

For the web-based visualization and interactivity we used ***Panel*** version 1.8.4 [20].

The web-application is served in the browser using the Bokeh Server Tornadocore application from ***Bokeh*** version 3.8.1 [21].

The initial bar-plots, presented for the APSS-based analyses, are visualized using ***hvPlot*** version 0.12.1 (https://hvplot.holoviz.org/en/docs/latest/index.html).

The APSS- and ANI-based distribution plots (jittered scatter and boxplot) visualizations, clustered heatmaps and multi-species boxplots are created using ***seaborn*** version 0.13.2 (https://doi.org/10.21105/joss.03021).

The network graph is visualized using ***hvPlot*** (cite) and ***NetworkX*** version 3.5 [25] packages. Saving network plots as images is done using the ***matplotlib*** (version 3.10.8) package [26].

The ANI vs. APSS scatter plot and the synteny per-position plots are visualized using the ***matplotlib*** version 3.10.8 package [26].

## Supplementary figures

**Figure S1:**
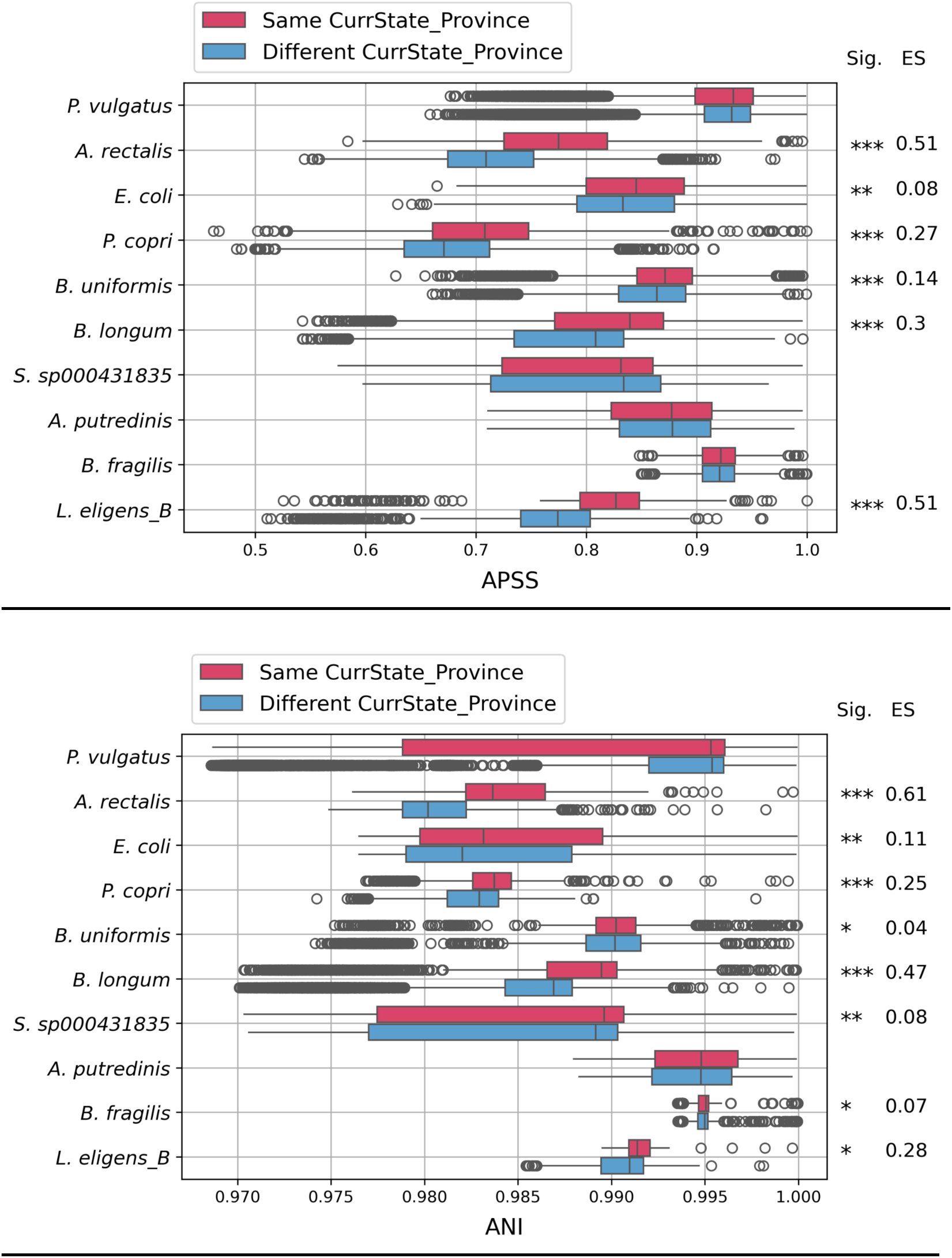
Multi-species analyses of strain-similarities of the most abundant 10 species in the human gut metagenomes collected in Gabon, Germany and Vietnam. Synteny-based (top) and ANI-based (bottom) comparisons, with internal within-species classification to sample pairs in which the hosts reside currently in the same province or not.

**Figure S2:**
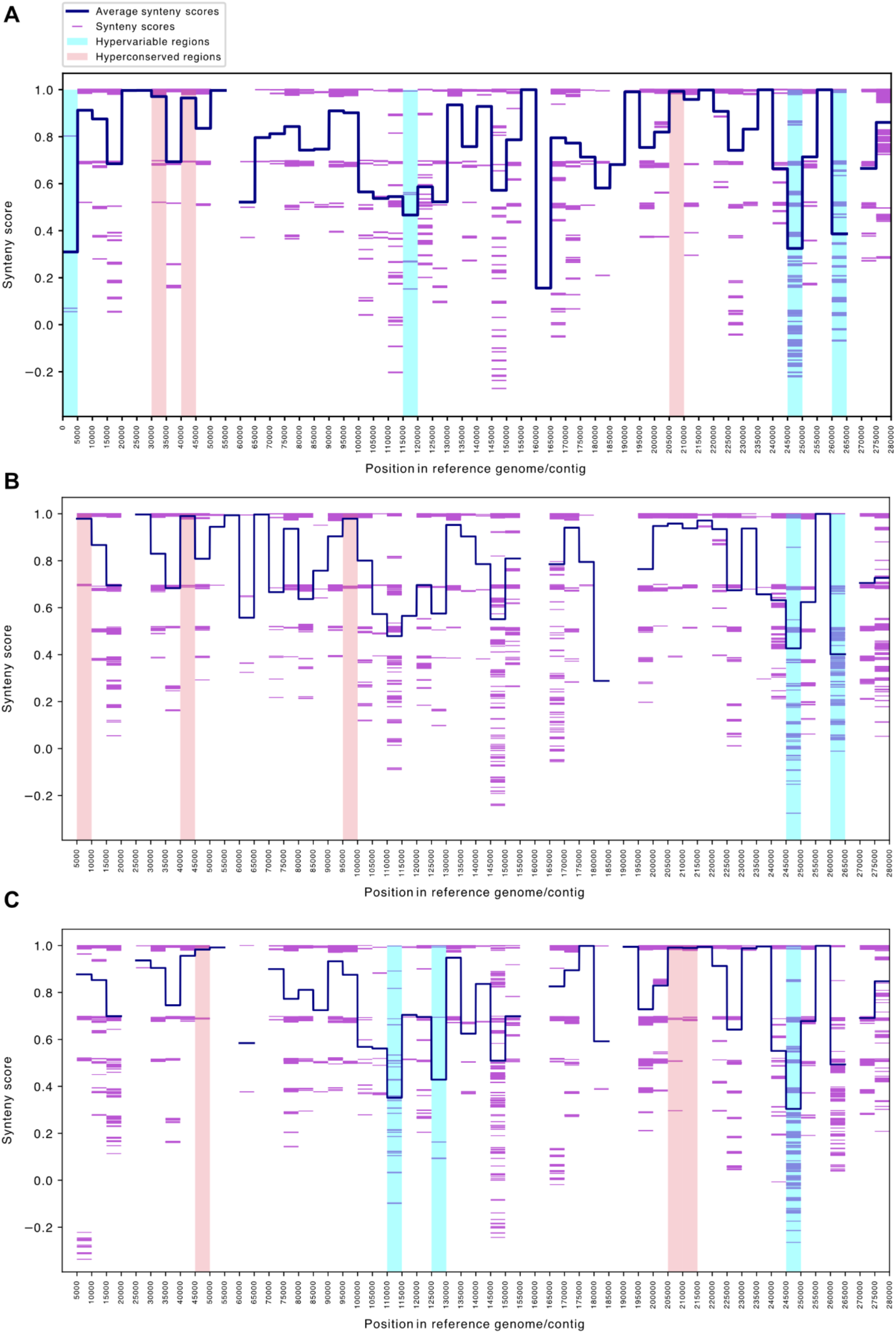
Synteny per-position plot of contig 630 of the *B.longum* reference genome, for each local population. Synteny per position plots for *B.longum* strains originating in (A) Germany (B) Gabon (C) Vietnam. Horizontal purple lines represent individual pairwise synteny scores for each region, the dark blue line represents the average synteny score for all pairwise comparisons, in each location. Vertical blue highlighting represents hypervariable regions, salmon vertical highlighting represents hyperconserved regions.

